# A chromosome-scale assembly of the major African malaria vector *Anopheles funestus*

**DOI:** 10.1101/492777

**Authors:** Jay Ghurye, Sergey Koren, Scott T. Small, Seth Redmond, Paul Howell, Adam M. Phillippy, Nora J. Besansky

**Affiliations:** Department of Computer Science, University of Maryland, College Park, MD; Genome Informatics Section, Computational and Statistical Genomics Branch, National Human Genome Research Institute, National Institute of Health, Bethesda, MD; Department of Biological Sciences, University of Notre Dame, South Bend, IN; Infectious Disease and Microbiome Program, Broad Institute, Cambridge, MA; Department of Immunology and Infectious Disease, Harvard TH Chan School of Public Health, Boston, MA; Centers for Disease Control, Atlanta, GA

## Abstract

**Background:** *Anopheles funestus* is one of the three most consequential and widespread vectors of human malaria in tropical Africa. However, the lack of a high-quality reference genome has hindered the association of phenotypic traits with their genetic basis in this important mosquito.

**Findings:** Here we present a new high-quality *An. funestus* reference genome (AfunF3) assembled using 240x coverage of long-read single-molecule sequencing for contigging, combined with 100x coverage of short-read Hi-C data for chromosome scaffolding. The assembled contigs total 446 Mbp of sequence and contain substantial duplication due to alternative alleles present in the sequenced pool of mosquitos from the FUMOZ colony. Using alignment and depth-of-coverage information, these contigs were deduplicated to a 211 Mbp primary assembly, which is closer to the expected haploid genome size of 250 Mbp. This primary assembly consists of 1,053 contigs organized into 3 chromosome-scale scaffolds with an N50 contig size of 632 kbp and an N50 scaffold size of 93.811 Mbp, representing a 100-fold improvement in continuity versus the current reference assembly, AfunF1.

**Conclusion:** This highly contiguous and complete *An. funestus* reference genome assembly will serve as an improved basis for future studies of genomic variation and organization in this important disease vector.

## Data Description

### Introduction and Background

Many insect genomes remain a challenge to assemble, and mosquito genomes have proven particularly difficult due to their repeat content and structurally dynamic genomes. These issues are compounded by the requirements of long-read sequencing technologies that typically require >10 μg of DNA for library construction. As a result, it is often impossible to construct a sequencing library from a single individual. Instead, sequencing a pool of individuals from an inbred population has been required [1]. For species that are amenable to extensive inbreeding, this approach has led to reference-grade genomes directly from the assembler [2]. However, when inbreeding is not possible, the sequenced pool of individuals can carry population variation that fragments the resulting assembly. In this case, instead of assembling a single genome, the assembler must reconstruct some unknown number of variant haplotypes.

Motivated by the goal of genome-enabled malaria control, a large international consortium previously sequenced and assembled the genomes of 16 *Anopheles* species using short-read Illumina sequencing [3,4]. Although these draft assemblies represented a crucial first step, their potential for 1) understanding and manipulating vectorial capacity traits, 2) inferring how key vector adaptations to hosts and habitats have arisen and are maintained, and 3) accurately defining vector breeding units and migration between them is constrained by two major limitations. First, many of these *Anopheles* assemblies are highly fragmented collections of relatively short scaffolds, causing gene annotation problems such as missing genes, missing exons, and genes split between scaffolds or sequencing gaps. Thus, one of the consequences of fragmented assemblies is that it is difficult to estimate gene copy number, which may be linked to important phenotypic traits (e.g. insecticide resistance) [5,6]. Genes of particular interest with respect to arthropod disease vectors (e.g., cytochrome P450s and odorant/gustatory receptors) may be especially prone to annotation errors, as many belong to gene families whose members are often physically clustered into tandem arrays.

A second major limitation of fragmented insect assemblies is that they are rarely scaffolded into chromosomes, owing to difficulty and lack of funding for physical or linkage mapping. Among other consequences, the unknown placement of scaffolds along chromosome arms means that their position within or outside of chromosomal inversions is difficult or impossible to determine. Many anopheline species are highly polymorphic for chromosomal inversions, which tend to occur disproportionately on particular chromosome arms [7–9]. In a heterozygote carrying one inverted and one uninverted chromosome, recombination between the reversed chromosomal segments is greatly reduced [10], creating cryptic population structure that can cause spurious associations in GWAS [11] and mislead recombination-based inference of selection and gene flow [12,13]. Importantly, chromosomal inversions also directly or indirectly influence traits affecting malaria transmission intensity—anopheline biting and resting behavior [14,15], seasonality [16], aridity tolerance [14,17–21], ecological plasticity [22,23] morphometric variation [24], and *Plasmodium* infection rates [25,26]. Thus, correct population genomic and GWAS inferences depend upon knowing the location of a marker in the genome.

*Anopheles funestus* is one of the three most important and widespread vectors of human malaria in tropical Africa [27–30], and unlike *Anopheles gambiae* with which it broadly co-occurs, it is a relatively neglected species. It is considered even more highly anthropophilic and endophilic than *An. gambiae* and amenable to conventional indoor-based vector control such as bed nets and indoor spraying of houses with residual insecticides. Indeed, historical house spraying campaigns in eastern and southern Africa not only locally eliminated this species, but the effect was maintained for several years following the cessation of spraying, due to the apparent inability of *An. funestus* to recolonize some areas. Likewise, *An. funestus* was eliminated from a humid forest and degraded forest areas in West Africa where malaria is meso- or hypoendemic [31]. However, in the savanna environment of West Africa where malaria is holo- or hyperendemic, similar historical indoor spraying campaigns failed to eliminate the species. Exophilic populations persisted which—despite marked anthropophily—continued to feed outdoors on cattle but also entered sprayed houses to bite humans. Today, the situation is worsened by the emergence and spread of insecticide resistance in this species [29,32–34].

Mastery over malaria will require tackling *An. funestus*, but it remains understudied; information on its behavior and genetics lags far behind *An. gambiae*. At least part of the reason for its neglect may be the historical lack of laboratory colonies, a problem solved with the establishment of the FUMOZ colony and its registration with the Anopheles program of BEI Resources (https://www.beiresources.org/AnophelesProgram.aspx). *An. funestus* shares with *An. gambiae* not only a broad sub-Saharan distribution and major vector status but also abundant chromosomal inversion polymorphism and shallow range-wide population structure [35]. However, there are behavioral and genetic heterogeneities relevant to malaria transmission that remain poorly understood. In West Africa, strong cytogenetic evidence points to cryptic, temporally stable assortatively mating populations co-occurring in the same villages [36–39]. These chromosomally recognized forms of *An. funestus*, named Kiribina and Folonzo, seem to differ in larval ecology and—importantly—they also differ in adult behaviors affecting vectorial capacity, most notably indoor resting behavior. Mechanistic understanding of the genomic determinants of these and other epidemiologically important phenotypic and behavioral traits ultimately depends on upgrading the *An. funestus* reference to a chromosome-based assembly in which the unanchored scaffolds are united, ordered and oriented on chromosome arms.

### Chromosome-scale assembly of *Anopheles funestus*

To achieve a complete and highly contiguous assembly of the *An. funestus* genome (AfunF3), we first assembled contigs from long, single-molecule reads, and then scaffolded these contigs into chromosome-scale scaffolds using Hi-C proximity ligation data. A similar strategy was recently used to improve the genome of *Aedes aegypti* [40]. An initial assembly of the long-read data alone (AfunF3 contigs) yielded a contig N50 size of 94.05 kbp (N50 such that 50% of assembled bases are in contigs of this size or greater) and extensive haplotype separation as evidenced by an inflated assembly size of 446.04 Mbp and a high rate of core gene duplications (48%) as measured by BUSCO [41]. These alternative alleles likely derive from natural variation circulating within the sequenced FUMOZ colony, as the DNA from a pool of adult mosquitoes was required for PacBio library preparation. Identifying and removing duplicate contigs via an all-vs-all alignment reduced the primary assembly size to 211.75 Mbp and improved the N50 size to 631.72 kbp (Table 1).

**Table 1:** Assembly statistics for the *An. funestus* genome. *AfunF1* represents the prior reference assembly, *AfunF3 contigs* denotes the complete long-read assembly with all contigs included and *AfunF3 primary* denotes the assembly after deduplication and scaffolding. QV(Illumina) denotes the assembly QV estimated using Illumina data and QV(10X) denotes the 10X Genomics data. QV(Illumina) is highest for the AfunF1 assembly, because it is the same data used to generate that assembly, whereas QV(10X) is based on data from a single mosquito of the same FUMOZ colony.

| Assembly | Number of Contigs | Contig N50 | Max Contig Size | Number of Scaffolds | Scaffold N50 | Max Scaffold size | Total Assembly Size | QV (Illumina) | QV (10X) |
| --- | --- | --- | --- | --- | --- | --- | --- | --- | --- |
| AfunF1 | 9,880 | 60,925 | 563,645 | 1,392 | 671,960 | 3,832,769 | 225,223,604 | 38.93 | 22.69 |
| AfunF3 contigs | 10,245 | 94,259 | 7,564,979 | 9,175 | 238,902 | 99,362,816 | 446,039,041 | 29.82 | 28.18 |
| AfunF3 primary | 1,053 | 631,722 | 7,564,979 | 3 | 93,811,348 | 99,362,816 | 210,827,327 | 24.94 | 25.82 |

The primary set of contigs (excluding alternative alleles) was then scaffolded using Hi-C Illumina reads to first bin the contigs into 3 chromosomes, followed by ordering and orientation of the contigs using the Proximo method (Phase Genomics, Seattle WA). The final scaffolded assembly (AfunF3 primary) contains 210.82 Mbp of sequence and a scaffold N50 of 93.81 Mbp. The resulting scaffolds represent the entirety of the three *An. funestus* chromosomes: 2, 3, and X (Figure 1).

**Figure 1:**
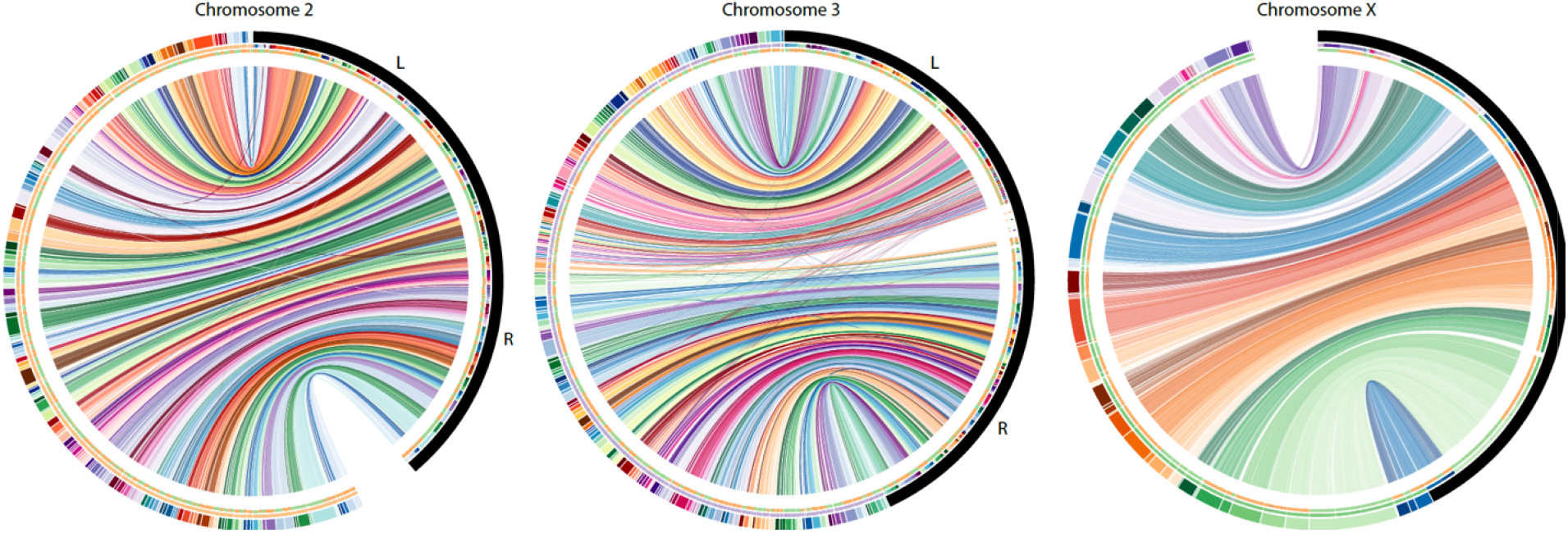
Circos plot comparing the AfunF1 assembly of *An. funestus* to the updated AfunF3 assembly. AfunF1 scaffolds (colored half of the outer ring) are ordered by majority alignment location onto AfunF3 (black half of the outer ring). Connecting lines indicate pairwise alignments between the two assemblies, and crossing lines indicate that part of the AfunF1 scaffold aligns to discordant regions on the AfunF3 chromosome. The first internal ring color correspond to the AfunF1 scaffold color. The second internal ring represents the orientation of the AfunF1 scaffolds onto AfunF3, where orange is forward and green is reverse.

Because single-molecule PacBio data is prone to insertion and deletion errors, all AfunF3 contigs were polished twice with Arrow [42] using the signal-level PacBio data and once with Pilon [43] using paired-end Illumina data from the same FUMOZ colony. Because Illumina-based polishing tools typically do not correct bases that appear heterozygous in the read set, we anticipated that variation in the FUMOZ colony would prevent the correction of variant bases. To help address this issue, we finally polished the assembly using 10X Genomics Illumina data obtained from an individual mosquito. As an independent test of base accuracy, we compared our new assembly (AfunF3 primary) and the prior assembly (AfunF1) to a 10X Genomics dataset from a different individual mosquito. The average Phred-scaled quality value [44] of the new assembly was estimated as QV28 versus QV23 for the Illumina-based AfunF1 assembly. This independent data indicates a higher average accuracy for the new assembly, but also revealed significant diversity within the colony. For example, calling variants using 10X Genomics data for two different mosquitos yielded widely different SNP counts (92,759 vs. 177,428).

We next evaluated the structural accuracy of the AfunF1 and AfunF3 assemblies by measuring their agreement with the raw PacBio reads. The intermediate assembly AfunF2 [45] was assembled before collection of all PacBio and Hi-C data, and so was deemed redundant and excluded from these analyses. When compared to the raw data, the AfunF3 primary assembly had fewer called structural differences (insertions, deletions, duplications, and inversions) than AfunF1 (Table 2). Despite the substantial single-nucleotide polymorphism observed within the FUMOZ colony, no large polymorphic inversions could be identified from the combined PacBio, Hi-C, and 10X Genomics data. Comparison of the chromosome-scale AfunF3 primary assembly versus the An. gambiae reference genome (AgamP4) confirmed a known reciprocal whole-arm translocation between 2L and 3R, as well as substantial intra-chromosomal shuffling (Figure 2). AfunF3 contigs also had fewer fragmented BUSCO core genes and a similar number of complete BUSCOs compared to AfunF1 (Table 2), but also a high rate of duplication. The AfunF3 primary scaffolds reduce duplication at the expense of lower BUSCO completeness.

**Table 2:** Validation of *An. funestus* genome assemblies using BUSCO gene set completeness, agreement of the assemblies with RNA-Seq transcriptome data, and structural accuracy inferred using PacBio long read data. *AfunF1* represents the prior reference assembly, *AfunF3 contigs* denotes the complete long-read assembly with all contigs included and *AfunF3 primary* denotes the assembly after deduplication and scaffolding. For BUSCO categories C denotes “Complete Genes”, S denotes “Single Copy Genes”, D denotes “Duplicated Genes”, F denotes “Fragmented Genes”, and M denotes “Missing Genes”. For long reads based structural variation, DEL denotes deletions, DUP denotes duplications, INV denotes inversions, and INS denotes insertions.

| Assembly | BUSCO statistics |  |  |  | Transcriptome data statistics |  |  | Structural variants called with long reads |  |  |  |
| --- | --- | --- | --- | --- | --- | --- | --- | --- | --- | --- | --- |
|  | C/S | C/D | F | M | Alignment Rate | Multi-mapped reads | % Transcripts in a single contig | DEL | DUP | INV | INS |
| AfunF1 | 2,756 | 16 | 27 | 16 | 81.79% | 23.92% | 84.96% | 9,036 | 455 | 152 | 3,798 |
| AfunF3 contigs | 2,765 | 1,068 | 18 | 17 | 84.34% | 36.97% | 91.16% | NA | NA | NA | NA |
| AfunF3 primary | 2,685 | 54 | 30 | 81 | 84.86% | 27.03% | 89.40% | 571 | 6 | 10 | 702 |

**Figure 2:**
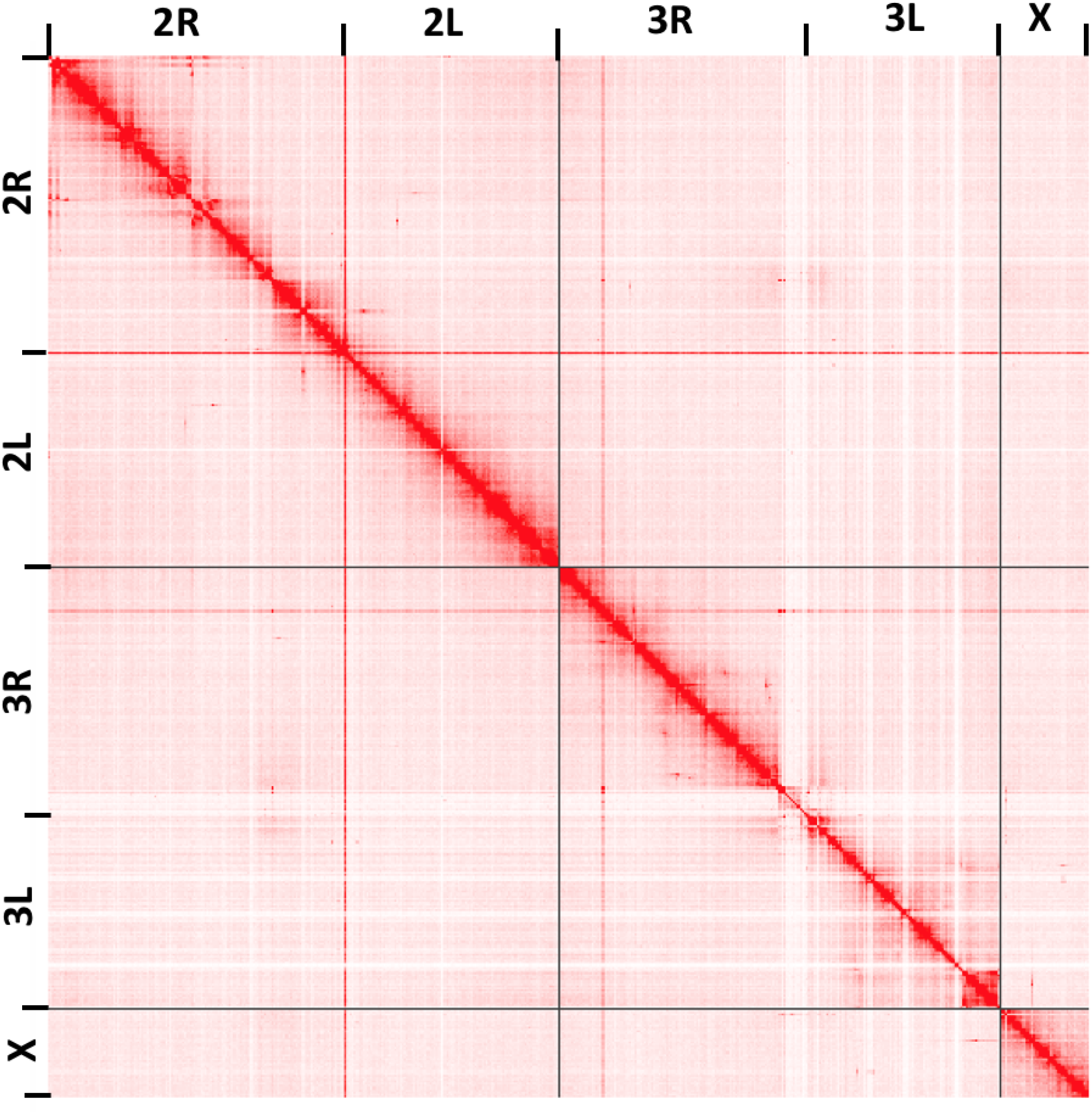
Hi-C interaction map for assembled *An. funestus* scaffolds generated using the Juicebox Hi-C visualization program [59]. Darker colors indicate a higher frequency of chromatin interaction. The plot shows clear separation of chromosome boundaries and limited off-diagonal interactions, supporting the global structure of the chromosome-scale scaffolds.

To further evaluate AfunF3’s suitability as an updated reference for *An. funestus*, we mapped RNA-Seq expression data to the assemblies and computed the number of concordant paired-end reads. A better assembly is expected to have both a higher fraction of mapped reads (completeness) as well as a higher fraction of correctly spaced and oriented pairs (structural accuracy). Both AfunF3 assemblies have better agreement of mapped read pairs as well as a higher overall mapping rate versus the AfunF1 assembly (Table 2). The AfunF3 contigs do have a higher rate of multi-mapping RNA-Seq reads, but this is reduced in the primary assembly while preserving the high mapping rate. In addition to a higher mapping rate, more complete transcripts were mapped to single contigs within the long-read assemblies. The average number of complete transcripts contained per contig was 67.38 for AfunF3 primary versus 5.28 for the AfunF1 assembly. These results demonstrate the greater continuity of the updated assembly, which provides sequence-resolved reconstructions of many *An. funestus* intergenic regions for the first time.

## Discussion

*Anopheles funestus* is one of the leading vectors of malaria and understanding the organization and function of its genome is key to controlling this deadly disease. Here we described a chromosome-scale assembly of the *An. funestus* genome using multiple sequencing technologies and assembly methods. The tremendous improvement in the completeness and contiguity of its genome will provide a valuable resource for future genomic analyses and functional characterization of this important species and enable a mechanistic understanding of the genomic determinants of epidemiologically important phenotypic and behavioral traits.

## Materials and Methods

### Library preparation and sequencing

A gravid female mosquito of the FUMOZ colony was allowed to lay eggs, and her offspring were inbred for a single generation. From this, an isofemale line was grown and DNA extracted from the adult females for sequencing with PacBio and Hi-C. 46 SMRT cells of PacBio RSII sequencing using the P6-C4 chemistry were run by the core facility at the Icahn School of Medicine at Mount Sinai (New York, NY), resulting in 173X coverage (assuming a 250 Mbp genome size). A previous study generated 70X coverage of the same colony using the older PacBio P5-C3 chemistry sequencing [45]. This older data was combined with the additional 173X coverage, totaling 60.95 Gb of long-read data in 10.93 million sequences (average length 5.6 kb, N50 read length 8.4 kb) and an estimated total coverage of 234X. Two Hi-C libraries were prepared and sequenced (one from mixed-sex larvae, the second from adult females) by Phase Genomics (Seattle, WA), resulting in ~100X coverage of Illumina Hi-C data containing ~187 million 80 bp paired-end Illumina reads.

### Assembly and scaffolding

PacBio contig assembly was performed with Canu v1.3 [46] using parameters: corOutCoverage=100 genomeSize=250m errorRate=0.013 batOptions=“-dg 3 -db 3 -dr 1 -ca 500 -cp 50”. The resulting contigs were then polished with Arrow [42] using default parameters and the P6-C4 PacBio signal data (because Arrow does not support the older P5-C3 data). After polishing, the assembly was separated into primary and alternative contigs to remove unnecessarily duplicated alleles from the AfunF3 contigs. This was performed using two different approaches. First, contigs containing at least one complete BUSCO gene were identified. For each BUSCO gene, if it was found contained in two or more contigs, the contig with the highest alignment score was kept as the primary. Next, all contigs not containing a BUSCO gene but assembled with high coverage (>40X) were added to the primary set.

To order and orient the primary contigs along the chromosomes, Hi-C reads were aligned using Bowtie2 [47] and scaffolding using Proximo (Phase Genomics, Seattle WA). Scaffold gaps spanned by PacBio reads were filled using PBJelly [48]. This assembly was again run through Arrow to polish the sequences inserted by PBJelly and fill any remaining short gaps. The Hi-C assembled scaffolds were then aligned using NUCmer [49] to the AfunF1 contigs for validation and the alignments visualized using Circos [50] and mummerplot. This identified a mis-join of chromosomes 3R and X, which was manually corrected. Additional manual curation using mapped transcripts, FISH probes [45], and comparison to AfunF1 scaffolds identified a few additional inversion errors in the scaffolds, mainly on distal 2L. Visual inspection of the Hi-C data showed clear signatures of scaffolding error. These errors were corrected by manually extracting the region and placing the sequence at the correct locus, as indicated by the Hi-C interactions. After these corrections, the scaffolded chromosomes (AfunF3 primary) show good agreement with the Hi-C data (Figure 3).

**Figure 3:**
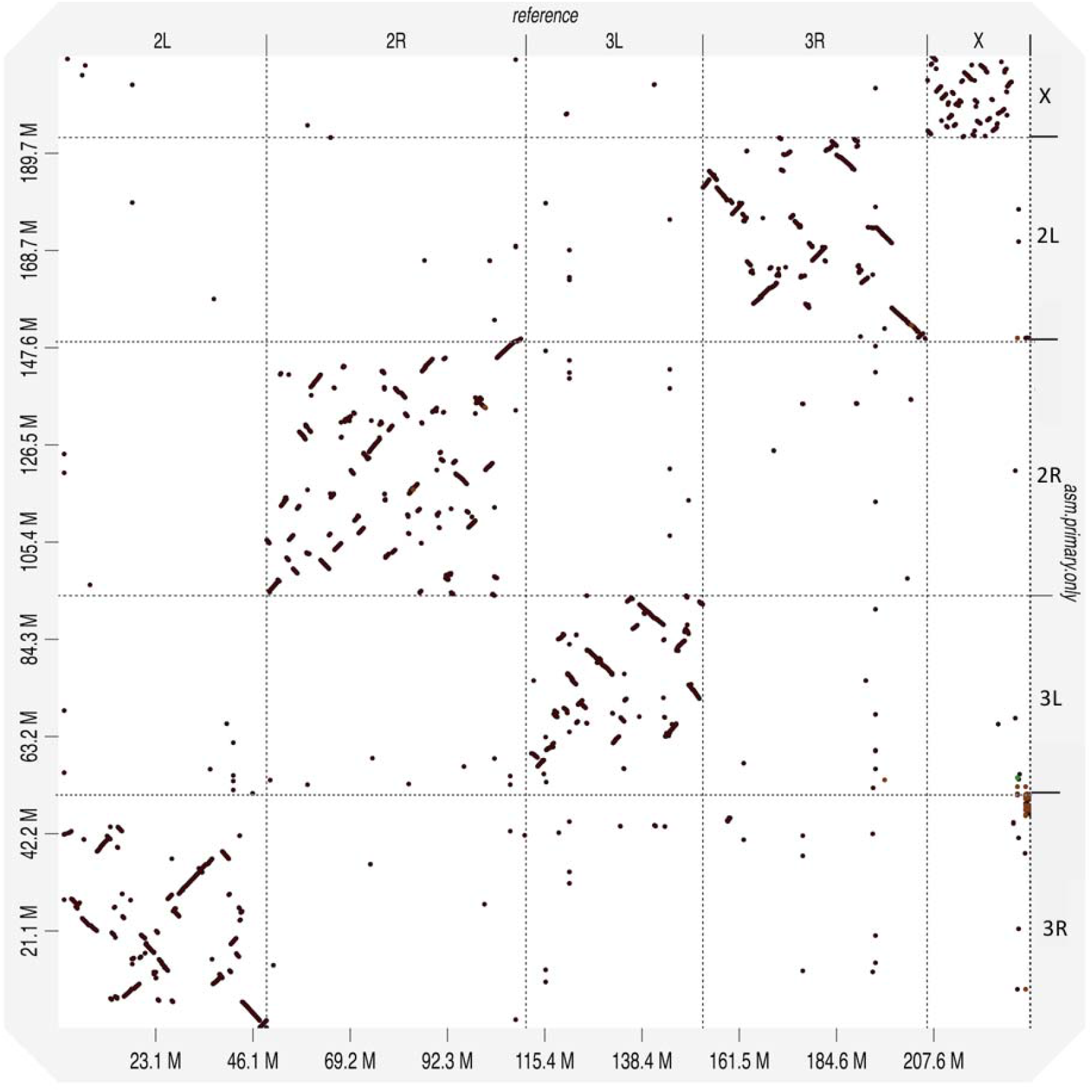
Whole genome alignment dotplot for *Anopheles funestus* and *Anopheles gambiae* genomes generated using D-GENIES [60]. A dot in the plot corresponds to a match between the corresponding genomic positions indicated on the axes. The *An. gambiae* reference genome is displayed on the x-axis, and the *An. funestus* AfunF3 primary assembly on the y-axis. A reciprocal whole-arm translocation between 2L and 3R is apparent, as well as substantial intra-chromosomal shuffling between these genomes.

As diploid and population variation introduces indels in the Arrow polishing process [51], the final assemblies were also polished by Pilon using paired-end Illumina data (NCBI SRA accession numbers: SRX209628 and SRX209387) and 10X Genomics Illumina data from a single individual (NCBI SRA accession number: SRX4819916). The paired-end Illumina data was mapped using BWA-MEM [52] and the 10X Genomics data mapped using Lariat [53] in a barcode-aware manner, as to improve the mapping quality. Consensus quality of the final assemblies was then estimated using an independent 10X Genomics dataset (NCBI SRA accession number: SRX4819903) of a different mosquito of the same FUMOZ colony. Based on the alignment of reads to the assembly, variants were called using freebayes (parameters: -C 2 -0 -O -q 20 -z 0.10 -E 0 -X -u -p 2 -F 0.5), and the assembly QV was estimated using called homozygous variants (i.e. positions where nearly all Illumina reads agreed with each other yet disagreed with the assembly).

### Validation

To check for the presence of contamination, assembled contigs were classified using Kraken [54] using a custom database including all microbial RefSeq genomes and all available mosquito genomes. Most of the assembled sequence (96.00%) was classified as *An. funestus* or Culicidae. The remaining sequences were primarily unannotated or annotated at a higher taxonomic level (3.76%), from possible bacterial/human sources (0.24%, 32 contigs), and had slightly lower GC content (Figure 4). However, none of these contigs were called contaminants by NCBI’s independent contamination check and so all contigs were included in the submitted assembly to avoid excluding novel mosquito sequence missing from the prior draft assemblies.

**Figure 4:**
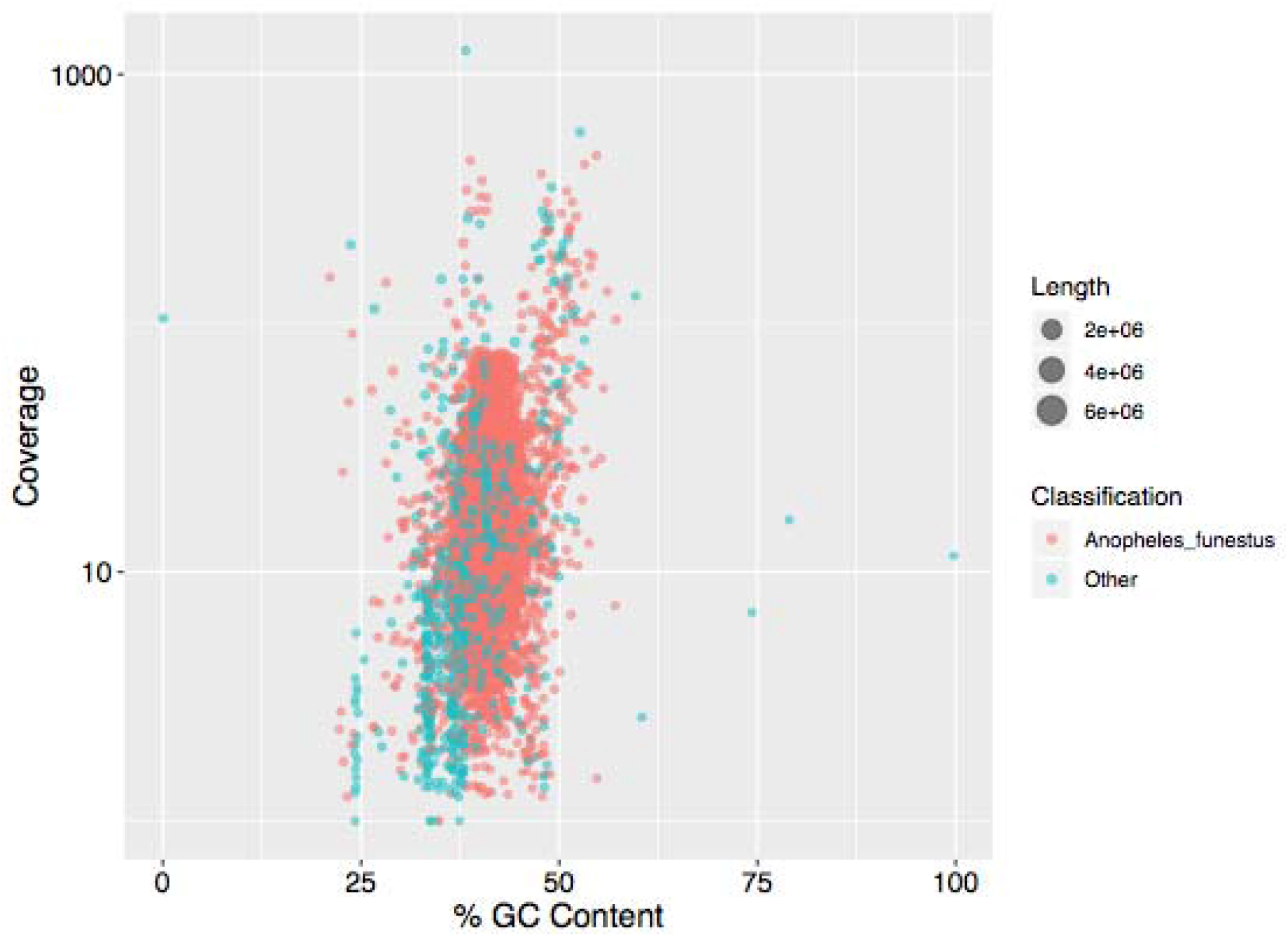
GC content versus coverage plot for all assembled *An. funestus* contigs. The orange points denote the contigs classified by Kraken as *An. funestus* and green points denote everything else. A majority of the contigs are classified as *An. funestus* by Kraken and there is no indication of extensive contamination.

The structural accuracy of the assemblies was evaluated by mapping raw PacBio reads and calling structural variants. PacBio reads were aligned to each assembly using NGMLR [55] with parameters: -t 16 -x pacbio --skip-write. Using these alignments, variants were called using Sniffles [55] with parameters: -t 32 -s 10 -f 0.25. Variants were then filtered to avoid capturing heterozygous population variants such that variants for which the alternate variant had ≥45 supporting reads and the assembly variant had <10 supporting reads were called as assembly errors.

Paired-end RNA-Seq for the *An. funestus* FUMOZ colony were downloaded from NCBI under accession SRR826832. These reads were aligned to all assemblies using the HISAT2 aligner [56] and assembled into transcripts using Trinity [57] with default parameters. The assembled transcripts were then mapped to all assemblies using GMAP [58]. Transcripts were required to be aligned over 90% of their length to a single contig to be considered “complete” in the assembly.

## Availability of supporting data

Raw genomic sequence reads are available in the NCBI Sequence Read Archive under project accession PRJNA494870. This Whole Genome Shotgun project has been deposited at DDBJ/ENA/GenBank under the accession RCWQ00000000. The version described in this paper is version RCWQ01000000.

## Declarations

### List of abbreviations

BUSCO: Benchmarking Universal Single-Copy Ortholog
PacBio: Pacific Biosciences
RNA-Seq: RNA-sequencing
NCBI: National Center for Biotechnology Information
SRA: Sequence Read Archive

### Ethics approval and consent to participate

Not applicable.

### Consent for publication

Not applicable.

### Authors’ contributions

AMP and NJB conceived and coordinated the project. JG, SK, STS, and AMP performed the genome assembly, validation, and comparative analyses. SR provided the 10X Genomics data and analysis. PH provided FUMOZ samples for sequencing. JG, AMP, and NJB drafted the manuscript. All the authors have read and approved the manuscript.

## Acknowledgments

The authors thank Ivan Liachko and Shawn Sullivan of Phase Genomics for assistance with Hi-C libraries and scaffolding, Robert Sebra of Mount Sinai for assistance with the PacBio sequencing, Igor Sharakhov of Virginia Tech for early access to the *An. funestus* FISH mapping data, and Rob Waterhouse of the University of Lausanne and Swiss Institute of Bioinformatics for assistance with Circos.

